# TUPAD: A Tn5-Mediated, Pre-amplification/Adapter-Dimer-Free Platform for Ultra-Uniform, One-Step PGT Profiling with High Stability and Broad Compatibility

**DOI:** 10.64898/2025.12.17.694820

**Authors:** Wenyang Yi, Kai Wu, Qiuyu Tang, Mengyao Wang, Hanxue Zhu, Huan Gao, Yudong Li, Mingyang Jiang, Hengzhi Wang, Yingjie An, Xiaomin Zheng, Qian Ouyang, Xia Zhao, Huaying Hu, Renqing Nie, Kai Wang, Duanru Cao, Yan Xu, Lijun Zhang, Tao Shu, Zhipeng Qu, Jiayin Guo, Lin Cao

**Author notes:** These authors contributed equally to this work (Wenyang Yi, Kai Wu, Qiuyu Tang, Mengyao Wang, Hanxue Zhu).

## Abstract

High-throughput sequencing stands as a cornerstone of clinical molecular diagnostics, particularly in preimplantation genetic testing (PGT)—a key application where the selection of chromosomally normal embryos is critical to enhancing in vitro fertilization (IVF) success rates. A fundamental challenge inherent to PGT, however, lies in the ultra-low gDNA input (typically <100 pg) from embryo biopsies. To mitigate this limitation, traditional workflows rely on whole genome amplification (WGA) technologies, such as MDA, PicoPlex, and MALBAC. Yet these WGA-based protocols inherently entail pre-amplification of gDNA prior to standard library construction-—a step that introduces significant coverage bias and compromised uniformity. These are intrinsic flaws irreparable by downstream library preparation or bioinformatics analysis, ultimately leading to clinical uncertainty in the interpretation of copy number variations (CNVs). In this study, we developed TUPAD (Tn5-Mediated, Uniform Pre-amplification-free and Adapter-Dimer-Free Library preparation), a transformative technology that bypasses the limitations of traditional WGA. By engineering a novel adapter-dimer-free transposon system, TUPAD enables high-efficiency library construction from as little as 1 picogram (pg) of gDNA. This breakthrough allows for a pre-amplification-free workflow, effectively eliminating the primary source of amplification bias. Compared to existing WGA standards, these advantages are fully retained in embryo biopsy and positive sample testing, TUPAD consistently generates highly uniform library data, significantly enhancing the detection limit and accuracy of CNVs and other genomic variants. Beyond these performance advantages, TUPAD also cuts wet-lab processing time by 50%. Practical validation confirms TUPAD results are stable, reliable, with proven practical utility in PGT testing.

## Introduction

Next-generation sequencing (NGS) has become a cornerstone technology for genetic variant detection in diagnostics, enabling rapid and accurate identification of pathogenic genomic mutations. Its advancement in reproductive medicine—particularly preimplantation genetic testing (PGT) for in vitro fertilized (IVF) embryos—facilitates the detection of chromosomal abnormalities, improving IVF success rates and preventing the transmission of genetic disorders^1,2^.

PGT involves biopsying 1 cell from a day-3 embryo or 5-10 cells from a day-5/6 blastocyst, followed by DNA sequencing to select chromosomally normal embryos for transfer^3–5^. Genomic variants detected in PGT are generally categorized as either structural chromosomal abnormalities or single-nucleotide variants (SNVs). Structural variants include insertions, deletions, duplications, inversions, and translocations, often resulting in copy number variations (CNVs) arising from misrepaired DNA double-strand breaks. Based on these variation types, PGT is classified into aneuploidy testing (PGT-A)^6,7^, monogenic disorder testing (PGT-M) ^8^, and testing for structural rearrangements (PGT-SR)(Fig. 1)^9,10^.

**Figure 1.**
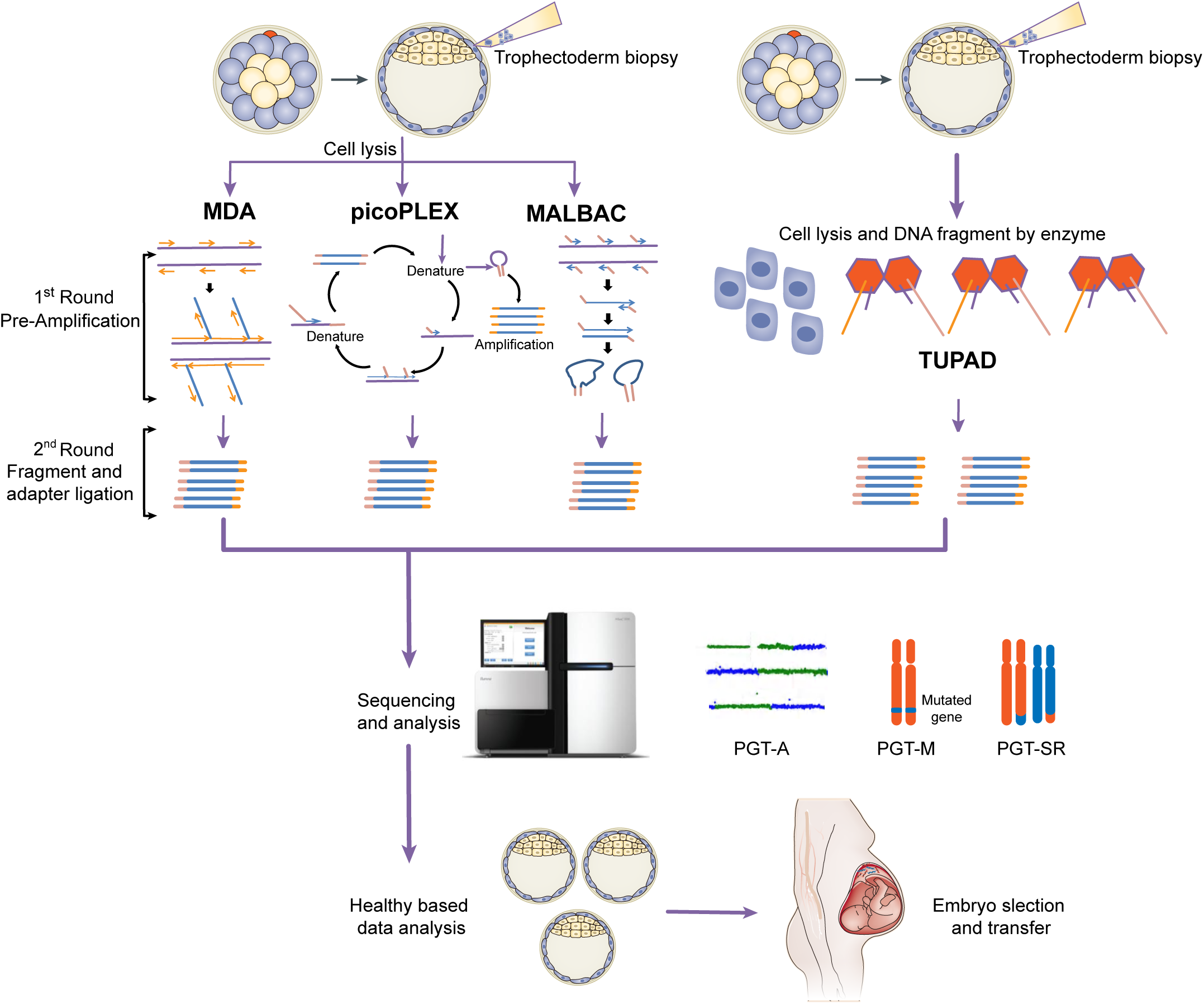
Schematic workflow of preimplantation genetic testing (PGT). The process begins with in vitro fertilization (IVF) to generate embryos, which are cultured to the blastocyst stage (Day 5-6 post-fertilization). Subsequently, trophectoderm biopsy is performed to collect a small number of cells (5-10 cells) from the outer layer of the blastocyst, preserving the inner cell mass to maintain embryonic developmental potential. Biopsied cells and subjected to genetic analysis, which may include techniques such as next-generation sequencing (NGS) for aneuploidy screening (PGT-A), targeted mutation analysis (PGT-M), or structural rearrangement detection (PGT-SR) based on clinical indications. After bioinformatics analysis and result validation, embryos identified as genetically normal or suitable for transfer are selected for subsequent embryo transfer procedures. In the optimized workflow, biopsied cells undergoes direct library construction using adapter-dimer-free Tn5 transpose, which enables unbiased whole-genome conversion without the need for conventional pre-amplification steps. This transposome-mediated library preparation allows simultaneous coverage of multiple genetic analyses, including next-generation sequencing based aneuploidy screening (PGT-A), targeted mutation analysis (PGT-M), and structural rearrangement detection (PGT-SR) according to clinical indications.

A primary challenge in PGT is the ultra-low DNA input (several to dozens of picograms) from single/trace cells, insufficient for traditional library preparation. All current PGT workflows thus rely on whole genome amplification (WGA) post-cell lysis, with strict contamination control ^6,7^. Successful WGA requires comprehensive coverage, uniform amplification, and high fidelity to avoid artifacts like allelic/locus dropout (ADO)—a key limitation exacerbated by the “Monte Carlo effect,” where low template abundance reduces the accuracy of genomic representation^11,12^. WGA also introduces sequence-dependent bias (e.g., GC content-related over/under-amplification) ^11^, chimeric amplicons (predominantly in multiple displacement amplification, MDA) ^13,14^, and non-template amplicons from contamination^14^. Failed WGA may necessitate re-sampling of precious embryonic tissue, potentially resulting in embryo loss.

A second critical limitation is the inability of existing WGA methods to simultaneously support PGT-A, PGT-M, and PGT-SR. Commonly used technologies each have inherent flaws: (1) Degenerate oligonucleotide-primed PCR (DOP-PCR) suits CNV detection but lacks coverage/uniformity for PGT-SR^8,9^; (2) Multiple displacement amplification (MDA) offers high coverage for PGT-M but exhibits inadequate uniformity and reproducibility for CNV analysis. As an MDA variant, PTA improves uniformity but requires specialized terminator bases and a terminator-tolerant phi29 DNA polymerase, yet yields short amplicons, is incompatible with PGT-SR, and lacks robustness to poor-quality samples^9,10,15–17^; (3) Multiple annealing and looping-based amplification cycles (MALBAC) and PicoPLEX enable reliable CNV/SNV detection but fail to amplify sequences with strong secondary structures, precluding PGT-SR ^11,15,18,19^. Due to the limited cell number in a single embryo sample, additional processing is required to enrich the genome^20,21^. Performing separate experiments is cumbersome, time-consuming, and increases costs. Therefore, overcoming the obstacles presented by multi-factor PGT parallel operations—by combining PGT-M, PGT-A, and PGT-SR into a single, universally applicable test—is highly clinically attractive.

Eliminating pre-amplification altogether and directly constructing sequencing libraries from lysed cells would fundamentally resolve amplification-induced bias, chimerism, and coverage distortion while preserving complete genomic information. However, conventional mechanical, enzymatic, or transposase-based library preparation protocols typically require nanogram(ng)-level DNA inputs and are incompatible with single-cell samples. At picogram input levels, we observed severe adapter dimer formation, which preferentially clusters during sequencing and drastically reduces usable data. Although we initially reduced adapter dimers in T4 ligase–based protocols through enzymatic digestion and bead-based size selection, such approaches proved incompatible with direct post-lysis library preparation due to the stringent buffer requirements of T4 ligase. While DOP-PCR enables pre-amplification-free trace-cells library construction, it exhibits poor uniformity due to random primer-mediated stochastic amplification^22^.

Transposase-based library construction has been shown to enable pre-amplification-free single-cell sequencing, but existing implementations depend on nanoliter-scale reactions and customized microfluidic platforms, limiting clinical scalability^23^. When we implemented Tn5-based library preparation at ultra-low inputs (1–10 pg) in standard reaction volumes (50 μL), consistently observed prominent non-mappable ∼144 bp fragments. Molecular characterization revealed these to be truncated adapter dimers generated during the tagmentation process. Strategies based on primer extension into Mosaic End sequences or phosphorothioate-modified primers partially reduced these artifacts but did not eliminate them. Complete suppression was achieved by employing a high-fidelity DNA polymerase lacking 3′–5′ exonuclease activity, preventing amplification of truncated adapter intermediates without reducing library yield. In summary, we have developed a refined Tn5-based, dimer-free technology that is fully compatible with 1 pg of gDNA input under standard reaction volumes, markedly enhancing sensitivity and robustness for ultra-low input DNA sample.

Based on this technology, we developed TUPAD (Tn5-Mediated, Uniform, Pre-amplification-free, and Adapter-Dimer-Free Library Preparation) — a next generation sequencing (NGS) platform integrating direct cell lysis and in situ genomic fragmentation. Engineered for ultra-low input scenarios, TUPAD enables direct single-cell library construction without pre-amplification, eliminating amplification artifacts and adapter dimers. Benchmarked against MDA, MALBAC, and PicoPLEX, TUPAD achieves superior genome coverage breadth, uniformity, and SNV/Indel detection accuracy in trace peripheral blood mononuclear cell (PBMC) samples, BRCA mutation standards, and embryo biopsy samples. Paired with a customized bioinformatics pipeline, it ensures 100% of embryo samples (N=85) meet diagnostic requirements while drastically reducing turnaround time and costs. TUPAD is poised to drive transformative advancements in PGT and related genetic testing.

## Results

### Eliminating T4 ligase-mediated adapter dimers via T7 endonuclease I and selective bead purification

For NGS library preparation, adapter dimer formation represents the key barrier to achieving successful ultra-low input library construction. Conventional DNA library preparation methods include mechanical, fragmentase, and transposase methods (Fig. S1A-B). Standard library preparation recommends 50-1000 ng input (low input mode down to 1 ng). Analysis of mechanical/fragmentase library products from 1 ng and 100 pg gDNA revealed increased adapter dimers with lower input, manifesting as abnormal peaks at 130-150 bp (Fig. S1C-D). Sanger sequencing confirmed that adapter dimers form via self-ligation of free adapter molecules in solution (Fig. S2A). Adapter dimers occupy sequencing flow cell space without useful data, usually removed by magnetic bead selection or agarose gel purification. Notably, magnetic bead selection is ineffective for low input dimer removal. Based on their formation mechanism, we developed two elimination methods: digestion and magnetic bead elution purification (Fig. 2A).

**Figure 2.**
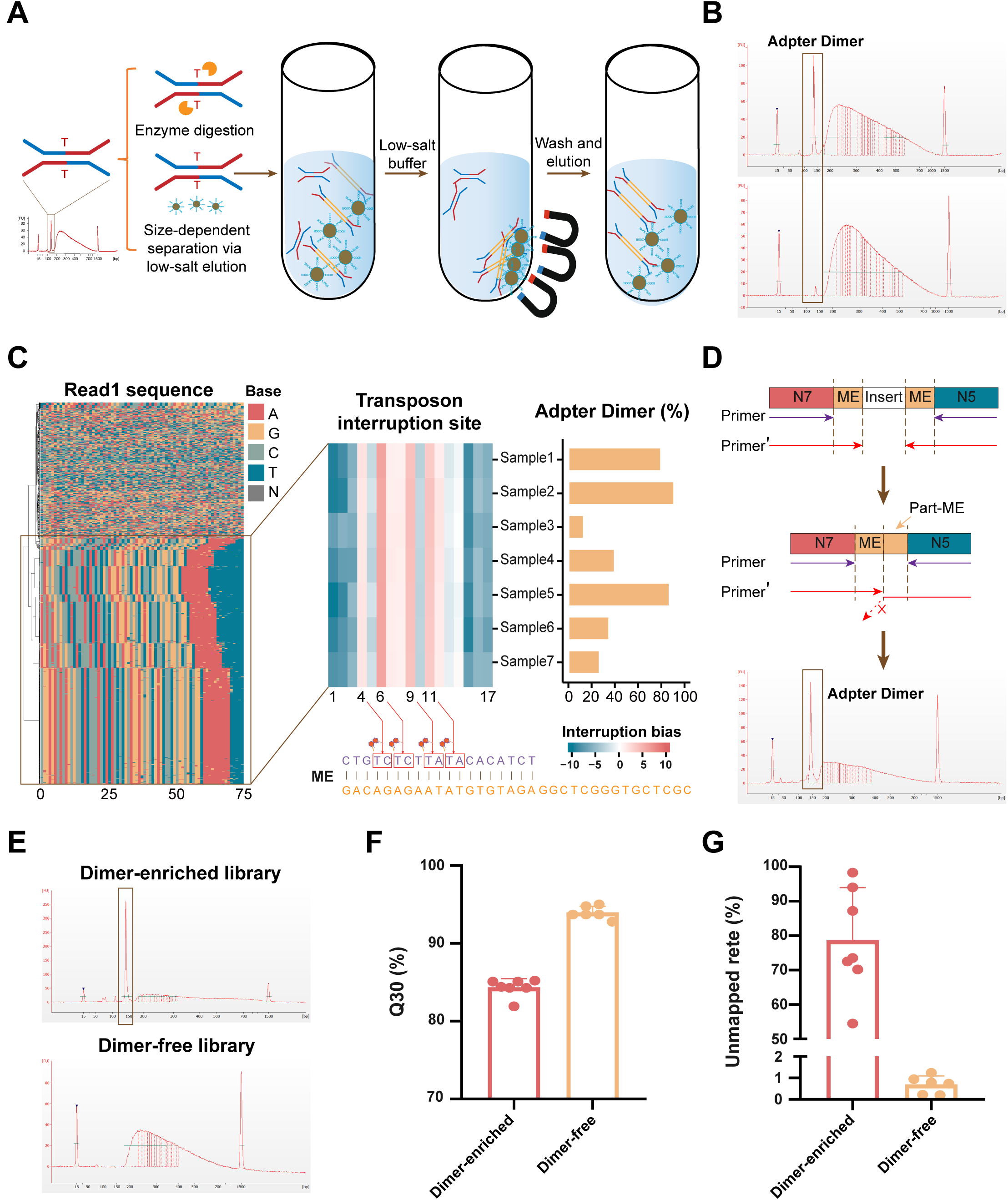
Adapter dimer formation in low-input library preparation and its elimination. (A) Schematic of adapter dimer removal strategies in ligation-based library preparation, including T:T mismatch endonuclease digestion and elution of adapter dimers during the bead purification step. (B) Size distribution of libraries analyzed by Agilent 2100 Bioanalyzer. The sharp peak at ∼130-150 bp corresponds to adapter dimers. The upper panel shows the library before 1U T7 endonuclease I with N001 buffer treatment, and the lower panel shows the library after 1U T7 endonuclease I with N001 buffer treatment. (C) Adapter dimer accumulation under low-input library preparation conditions. The left heatmap panel shows the base composition of 1,000 randomly sampled R1 reads from a representative library. The brown box highlights R1 sequences corresponding to adapter dimers. Across seven representative libraries, Tn5 preferentially interrupts the mosaic end (ME) sequence at positions 4, 6, 9, and 11, indicating sequence bias toward TC or TA motifs. The right panel summarizes the proportion of dimer reads in these seven libraries, which ranges from 10% to 90%. (D) The upper panel represents a library with a normal structure, characterized by intact N5 and N7 indices flanking a complete mosaic end (ME) sequence, resulting in sequencing reads corresponding to properly inserted DNA fragments. The lower panel shows the presence of partial ME (part-ME) sequences, which arise from Tn5-mediated fragmentation of the ME sequence and reflect aberrant library structures. These libraries are generated by conventional primer (purple arrow) amplification. The extension primer (red arrow) will inhibit the abnormal libraries from being enriched through amplification. Size distribution of libraries analyzed by Agilent 2100 Bioanalyzer, shows the library amplified with extension primer. (E) Size distribution of libraries analyzed by Agilent 2100 Bioanalyzer. The sharp peak at ∼144 bp corresponds to adapter dimers. The upper panel shows the dimer-enriched library before Tn5 adapter-dimer-free technology treatment, and the lower panel shows the dimer-free library treated with Tn5 adapter-dimer-free technology. (F) Q30 statistics of sequencing data for libraries with and without dimer sequences. Q30 values were calculated from clean reads after adapter trimming. Because part-ME sequences are truncated at the 5’ end, standard adapter-trimming parameters fail to remove these fragments, leading to a reduction in Q30 in the processed reads. (G) Unmapped read fraction for libraries with and without dimer sequences. As dimer sequences are not effectively removed by standard adapter-trimming, they enter the mapping process and result in a substantial increase in the proportion of unmapped reads.

Since adapter dimers arise from adapter self-ligation, we used T7 endonuclease I (mismatch-specific) for digestion. Spiking synthetic annealed adapter dimers into libraries, we observed improved dimer removal with increasing enzyme dosage. At 0.5U enzyme, 64.3-83.8% of dimers were removed compared to controls (CTR), 2U consistently achieved >90% dimer removal with ∼20.7-32.8% library loss (Fig. S2B). Otherwise, it did not impact sequencing quality or mutation detection in NA12878 gDNA samples (figureS2 C-D). Poor magnetic bead selection efficiency stems from the solid phase reversible immobilization (SPRI) principle: DNA condenses from linear to globular at PEG concentrations above the critical point, precipitating and reverting to a helical conformation below it^24^. Independent of PEG concentration, the critical point correlates with DNA size, decreasing as fragment length increases^25^. Accordingly, we hypothesized that library molecules bind tightly to beads at high PEG/salt concentrations, whereas short adapter dimers crowd between library DNA and beads in low-input samples. Following initial bead incubation, resuspension in clean buffer adjusts PEG/salt concentrations to maintain library condensation (loose bead binding) while releasing free dimers into solution. Based on this principle, we developed a low-salt, low-PEG buffer (Vazyme N001) that removed 52.9-65.8% of adapter dimers with <10% library loss (Fig.S2 E-F).

Combining endonuclease digestion and magnetic bead elution, we optimized 1U T7 endonuclease I with N001 buffer, which achieved near-complete adapter dimer removal for 1 pg input in standard and fragmentase-based library preparation with minimal library loss (Fig.2B).

### Tn5-based library preparation at ultra-low DNA input

The transposase-adapter complex enables one-step DNA fragmentation and adapter ligation, shortening the experimental workflow with a minimum input of 1 ng DNA. At ultra-low inputs, many short fragments around 144 bp appear (Fig. S1E).

To identify these impurities, we analyzed sequencing data and found fragments unmappable to the genome, not matching complete adapter sequences, These were adapter dimers formed after transposon interruption, harboring TA/TC motifs and accounting for 10–90% of clean reads (Fig. 2C-D). Cloning and sequencing of these short fragments confirmed they were indeed broken adapter dimers (Fig. S3A). To address this issue, we targeted blocking adapter dimer PCR enrichment to ensure library purity. Given the dimers’ sequence features, we hypothesized that adapter nucleotide extensions would disrupt primer binding sites, abrogating dimer enrichment. Using engineered extended adapters, sequencing data showed ∼144 bp short fragments persisted—this initial strategy was ineffective (Fig. 2D). This failure likely arises from the 3′-5′ exonuclease activity of high-fidelity DNA polymerases, which trims extended primer regions and restores primer binding to adapter dimers. Based on this hypothesis, we modified 1-3 terminal primer nucleotides with phosphorothioate (resistant to exonuclease cleavage), markedly reducing adapter dimers (Fig. S3B). However, phosphorothioate modifications showed limited efficacy, prompting us to switch to a high-fidelity DNA polymerase lacking 3′-5′ exonuclease activity. This approach completely eliminated adapter dimers without compromising library yield, establishing a robust Tn5 adapter-dimer-free library protocol. With this optimized Tn5 library technology, we next evaluated its performance for ultra-low input samples (≤1 ng gDNA), which was compatible down to 1-10 pg—successfully constructing libraries with complete adapter dimer elimination (Fig. 2E). In terms of sequencing quality, the Tn5 adapter-dimer-free technology significantly improved Q30 scores and markedly reduced unmapped reads (Fig. 2F-G), enhancing library success rates and lowering costs.

In summary, we show that short fragment impurities in ultra-low input transposase libraries arise from transposase-induced adapter dimers. We developed an innovative Tn5 adapter-dimer-free technology that efficiently eliminates these dimers, abolishing their interference with usable sequencing data and boosting experimental success. This offers great potential for ultra-low input library preparation.

### TUPAD (Tn5-Mediated, Uniform Pre-amplification-free and Adapter-Dimer-Free Library preparation)

While both T4 ligase and Tn5 transposase enable adapter-dimer-free library construction, T4 ligase-based protocols are unsuitable for direct single-cell library prep due to two critical limitations: first, it lacks intrinsic genomic DNA fragmentation activity, necessitating a separate fragmentation step; second, it depends on purified, high-integrity DNA templates and is highly sensitive to cell lysis buffer inhibitors (e.g., residual proteases, chaotropes), which severely impair ligation efficiency. In contrast, Tn5 transposase facilitates one-step fragmentation and adapter ligation, exhibits stable activity in mild lysis buffers, and requires no prior DNA purification, enabling direct single-cell library construction.

TUPAD was developed by integrating the Tn5 adapter-dimer-free system with a robust cell lysis buffer, enabling direct library construction. It is ideally suited for single-cell and low-input samples, with significant advantages particularly in PGT. We compared the performance of uniformity and mutation detection with that of three commercial single-cell whole-genome amplification protocol (QIAGEN-MDA, Yikon-MALBAC, TAKARA-PicoPLEX) on trace cell samples in 1-6 PBMC cell. To assess genome uniformity, multiple statistical metrics were employed, revealing that TUPAD exhibited superior genome coverage (Fig. 3A), the lowest median of absolute pairwise differences (MAPD; Fig. 3B, Fig. S4A), and the lowest coefficient of variation (CV) for genome coverage (Fig. 3C, Fig. S4B)—even at the single-cell level. Raw read density across all 23 chromosomes reveals that TUPAD exhibits the most uniform CNV profiles among the four methods—even without normalization—outperforming PicoPLEX (Fig. 3D).

**Figure 3.**
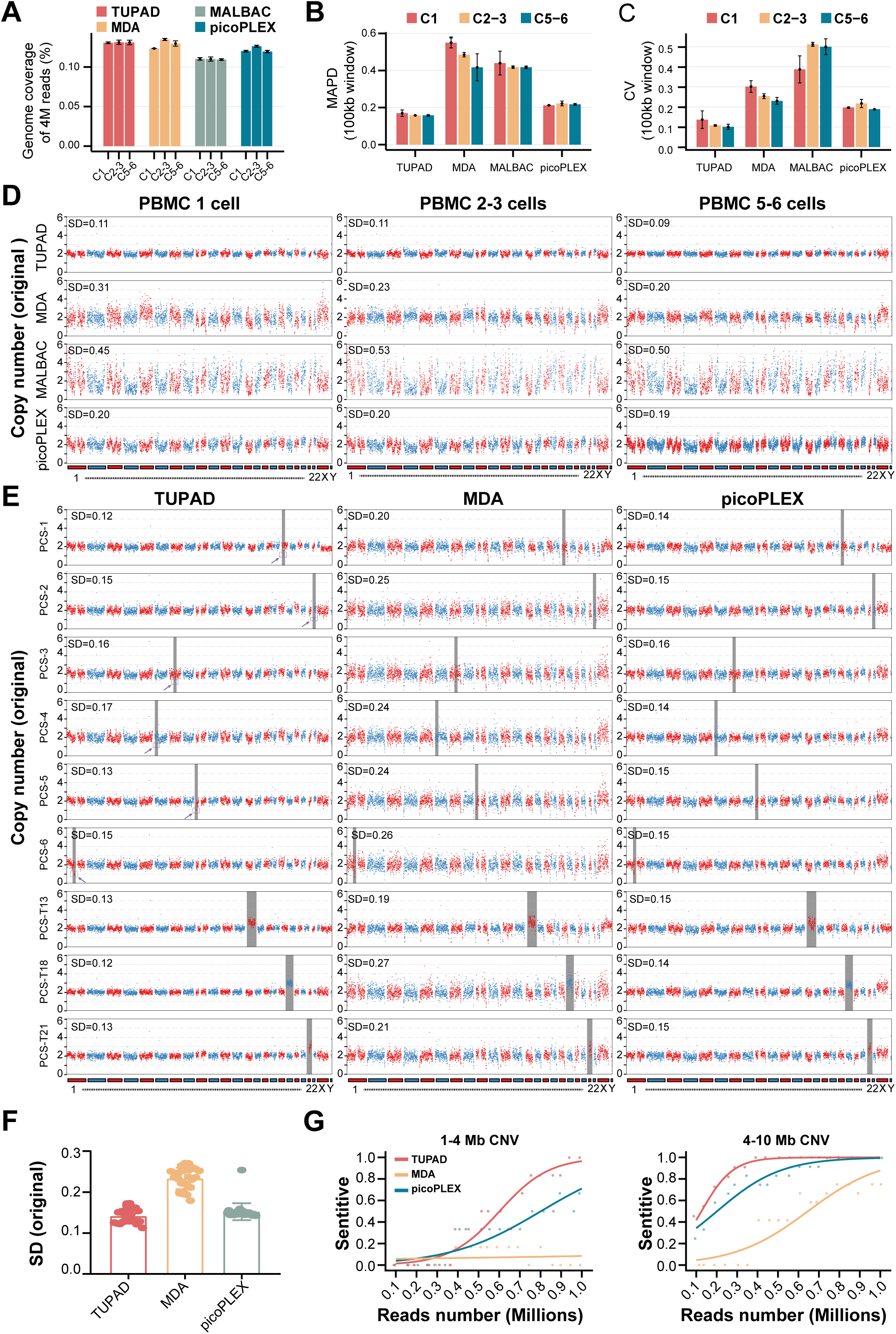
Comparison of coverage uniformity and CNV detection among TUPAD and other WGA methods. (A) Genome coverage across different methods with increasing numbers of input cells. C1, C2-3, and C5-6 denote libraries prepared from 1 cell, 2-3 cells, and 5-6 cells, respectively. All methods were performed in duplicate, and sequencing data were downsampled to 4 million reads per sample (∼0.2× genome sequencing depth). (B) Genome coverage uniformity measured by the median absolute pairwise difference (MAPD) across increasing cell numbers (1, 2-3, or 5-6 cells). MAPD was calculated using 100 kb genomic bins, with two technical replicates per method. (C) Genome coverage uniformity measured by the coefficient of variation (CV) across increasing cell numbers (1, 2-3, or 5-6 cells), using 100 kb genomic bins and two technical replicates per method. (D) Original copy-number profiles for increasing numbers of peripheral blood mononuclear cells (PBMCs) generated using TUPAD, MDA, MALBAC, and picoPLEX. Standard deviation (SD) was calculated from raw read counts in 600 kb windows. TUPAD consistently exhibited lower SD and more stable copy-number profiles across all input cell numbers (1, 2-3, or 5-6 cells). (E)Original copy-number profiles of 9 PGT-A positive control standards (PCS) generated by TUPAD, MDA, and picoPLEX. PCS1-6 correspond to CNV deletions <10 Mb, whereas PCS13, 18, and 21 represent whole-chromosome trisomies. Dark gray boxes indicate chromosomes harboring CNVs; purple boxes and arrows highlight smaller CNV deletions uniquely detected by TUPAD. (F) Boxplots showing SD statistics for the copy-number profiles of the 9 positive control standards (corresponding to panels in D) across TUPAD, MDA, and picoPLEX. (G) CNV detection performance for positive reference standards at low sequencing depths (0.1-1 million reads). The left panel shows detection sensitivity for CNVs of 1-4 Mb (two CNVs, three replicates, six CNV events in total), and the right panel shows detection sensitivity for CNVs of 4-10 Mb (four CNVs, three replicates, twelve CNV events in total). For each data point, the x-axis indicates the method at a given read depth, and the y-axis represents CNV detection sensitivity (criteria described in Methods). Curves were fitted using a softmax regression. TUPAD exhibited higher sensitivity for detecting smaller CNVs.

Leveraging its excellent coverage and uniformity in low-input cell samples, TUPAD’s reliability for PGT-A variant detection was evaluated using PGT-A positive control standards harboring diverse genomic variants. Given MALBAC’s inferior uniformity compared to other WGA methods (Fig. 3A–D, Fig. S4A–B)^26^, MDA and PicoPLEX were included as comparators in subsequent experiments. Raw read density profiles across all 23 chromosomes demonstrated that all three methods effectively detected whole-chromosome abnormalities—including common aneuploidies involving chromosomes 13, 18, and 21—whereas TUPAD exhibited more distinct and reproducible signal alterations for smaller CNVs (<10 Mb), underscoring enhanced resolution at the sub-chromosomal level (Fig. 3E), with uniformity comparable to PicoPLEX and superior to MDA (Fig. 3F). To further compare the detection sensitivity of the three methods under low sequencing depth, we subsampled the data to no more than 1 million reads and evaluated the sensitivity for detecting small CNVs(<10Mb). Under these conditions, TUPAD consistently demonstrated superior detection performance compared with the other methods (Fig. 3G), attesting to its superior sensitivity at low sequencing depths and thereby translating to cost savings in practical clinical testing.

Collectively, compared to traditional WGA methods, TUPAD exhibits superior uniformity and coverage in single-cell sequencing, enabling accurate detection of genomic variants across different size ranges in PGT-A positive standards. With higher sensitivity at lower sequencing depths, it underscores significant potential for practical PGT-A applications.

### TUPAD achieves superior mutation detection performance

Beyond structural variations, many monogenic diseases arise from single-gene mutations, imposing stringent demands on sequencing coverage and fidelity in PGT-M applications^27^. To assess TUPAD’s capacity to preserve genomic integrity, we evaluated its mutation detection performance with a standard BRCA genomic DNA (gDNA) reference. For the control, 1 ng of BRCA gDNA was used to generate whole-genome sequencing (WGS) libraries via a commercial Tn5-based library preparation protocol.

All samples were processed using the same pipeline and downsampled to an equal number of reads, approximately 300 million single-end reads were obtained per sample, corresponding to ∼15× whole-genome sequencing depths. Genome coverage uniformity was evaluated using multiple complementary metrics (see methods).

TUPAD consistently achieved higher genome coverage breadth than MDA and picoPLEX across increasing sequencing depths and even slightly exceeded the WGS input below 140 million reads (Fig. 4A). Lorenz curve analysis showed that TUPAD closely recapitulated the coverage uniformity of bulk WGS and outperformed other WGA methods (Fig. 4B), which was further supported by lower Gini index values (Fig. 4C). Consistent results were obtained from MAPD and CV analyses, as well as genome-wide copy-number profiles, indicating highly uniform and high-coverage libraries generated by TUPAD (Fig. S5A–B).

**Figure 4.**
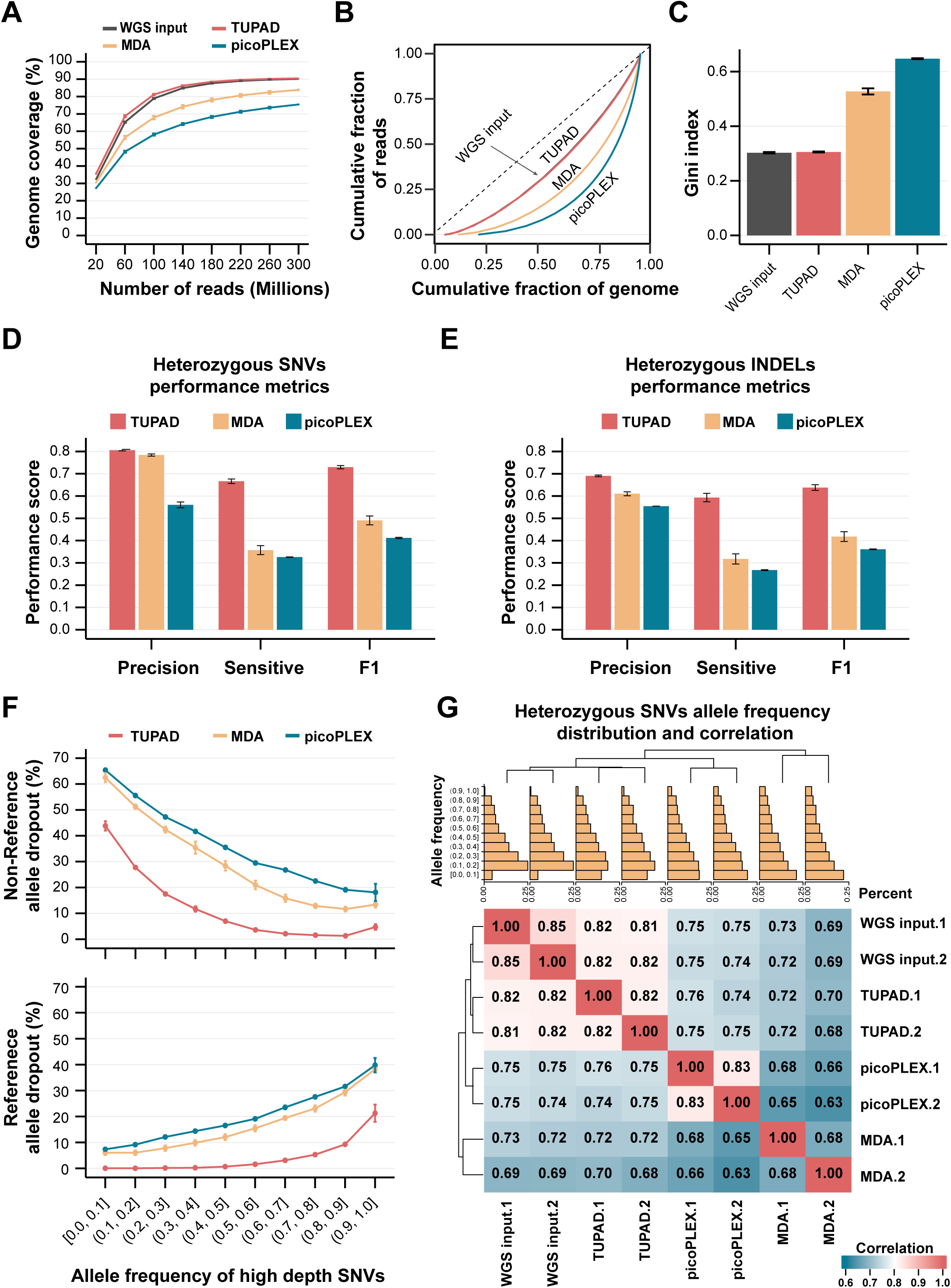
TUPAD improves genome coverage breadth and variant detection performance. (A) Genome coverage breadth across different methods as a function of increasing numbers of single-end sequencing reads, measured by the fraction of the genome covered. TUPAD showed coverage breadth comparable to WGS input across all sequencing depths; 300 million 150-bp single-end reads correspond to approximately 15× whole-genome coverage. (B) Genome coverage uniformity across methods illustrated by Lorenz curves, where the diagonal line represents perfectly uniform coverage and increasing deviation reflects greater coverage bias. TUPAD most closely recapitulated the coverage uniformity observed in WGS input. (C) Genome coverage uniformity quantified by the Gini index across different methods. Lower Gini index values indicate more uniform coverage, and TUPAD exhibited low and highly reproducible Gini index values comparable to bulk DNA and substantially lower than those of other WGA methods. (D) Detection performance of heterozygous SNVs across different allele-frequency bins for TUPAD and three WGA methods, evaluated using precision, sensitivity, and F1 score. TUPAD exhibited improved overall SNV detection performance compared with other WGA methods. (E) Detection performance of heterozygous INDELs across different allele-frequency bins for TUPAD and three WGA methods, evaluated using precision, sensitivity, and F1 score.TUPAD consistently outperformed other WGA methods in INDEL detection accuracy. (F) Non-reference allele dropout (NR-ADO) rates of heterozygous SNVs stratified by allele frequency, calculated from sites with ≥20× coverage in both replicated bulk WGS inputs. TUPAD showed reduced NR-ADO rates across allele frequency ranges compared with other WGA methods. (G) Reference allele dropout (Ref-ADO) rates of heterozygous SNVs stratified by allele frequency, calculated from sites with ≥20× coverage in both replicated bulk WGS inputs. TUPAD exhibited lower Ref-ADO rates, indicating improved allelic balance. (H) Correlation of heterozygous SNV allele frequencies among TUPAD, bulk WGS input, and three WGA methods, visualized by a correlation heatmap. Only SNVs with ≥20× sequencing depth across all samples were included (7,406 SNVs), and TUPAD showed the highest Pearson correlation with bulk WGS input; bar plots above the heatmap show the allele frequency distributions detected by each methods.

To evaluate mutation detection performance, variants called by each method were compared with those detected in the WGS input and classified as true positives (TP), false positives (FP), or false negatives (FN), which were subsequently used to calculate precision, sensitivity, and F1 score (see methods). TUPAD exhibited superior SNV detection performance, particularly in terms of sensitivity and F1 score, compared with other WGA methods (Fig. 4D). Notably, this advantage was robust across different truth-set depth thresholds derived from the WGS input (Fig. S6A). A similar improvement was observed for INDEL detection (Fig. 4E, Fig. S6B). For allelic amplification fidelity analysis, allele dropout rates were quantified at heterozygous variants defined in the WGS reference. Given that the standard BRCA gDNA reference does not always exhibit mutation allele frequencies centered at 0.5, heterozygous SNVs were stratified by their truth-set allele frequencies. TUPAD showed significantly reduced allele dropout rates relative to MDA and PicoPLEX (Fig. 4F).

Allelic skewing in mutation detection was further evaluated by examining depth distributions of TP heterozygous variants, which revealed TUPAD patterns closely resembling those of WGS input (Fig. S7A-B). Consistently, mutation spectrum bias analysis demonstrated minimal deviation of TUPAD from the WGS input (Fig. S8A-B). Given the modest effective depth (∼15×) across samples, only heterozygous SNVs with high coverage (≥20×) were included in allele frequency correlation analysis. TUPAD exhibited a high pearson correlation (>0.8) with the WGS reference and a more comparable allele frequency distribution than other WGA methods (Fig. 4G).

Overall, compared with widely employed WGA methods, TUPAD exhibits substantially improved accuracy and sensitivity for SNV and INDEL detection in low-input genomes. Its high-fidelity amplification minimizes allelic dropout and skewing, effectively suppresses amplification-induced artifacts, and underscores strong potential for practical PGT-M applications.

### TUPAD Delivers Superior practical PGT Performance via Integrated Algorithms

TUPAD’s high coverage, uniformity, and fidelity in single-cell library preparation underscore its advantage for practical PGT detection within a consistent workflow. In practical testing using clinical embryo samples, TUPAD was comprehensively compared with conventional WGA methods. Raw read density across all 23 chromosomes demonstrated that TUPAD’s uniformity was superior to PicoPLEX and MDA (Fig. 5A). Despite exhibiting favorable uniformity in these clinical embryo samples, its performance was inferior to that observed in gDNA and single-cell samples (Fig. 3A-E)—a phenomenon attributed to abundant heterochromatic regions in early-stage embryonic cells, which impede direct Tn5-mediated fragmentation^28^. We present an improved CNV detection algorithm that establishes a baseline via outlier removal and GC bias correction. To enhance accuracy, the algorithm further applies a standard deviation (SD) correction derived from a background cohort to windows with significant sequencing variability (Fig. 5B; see Methods).

**Figure 5.**
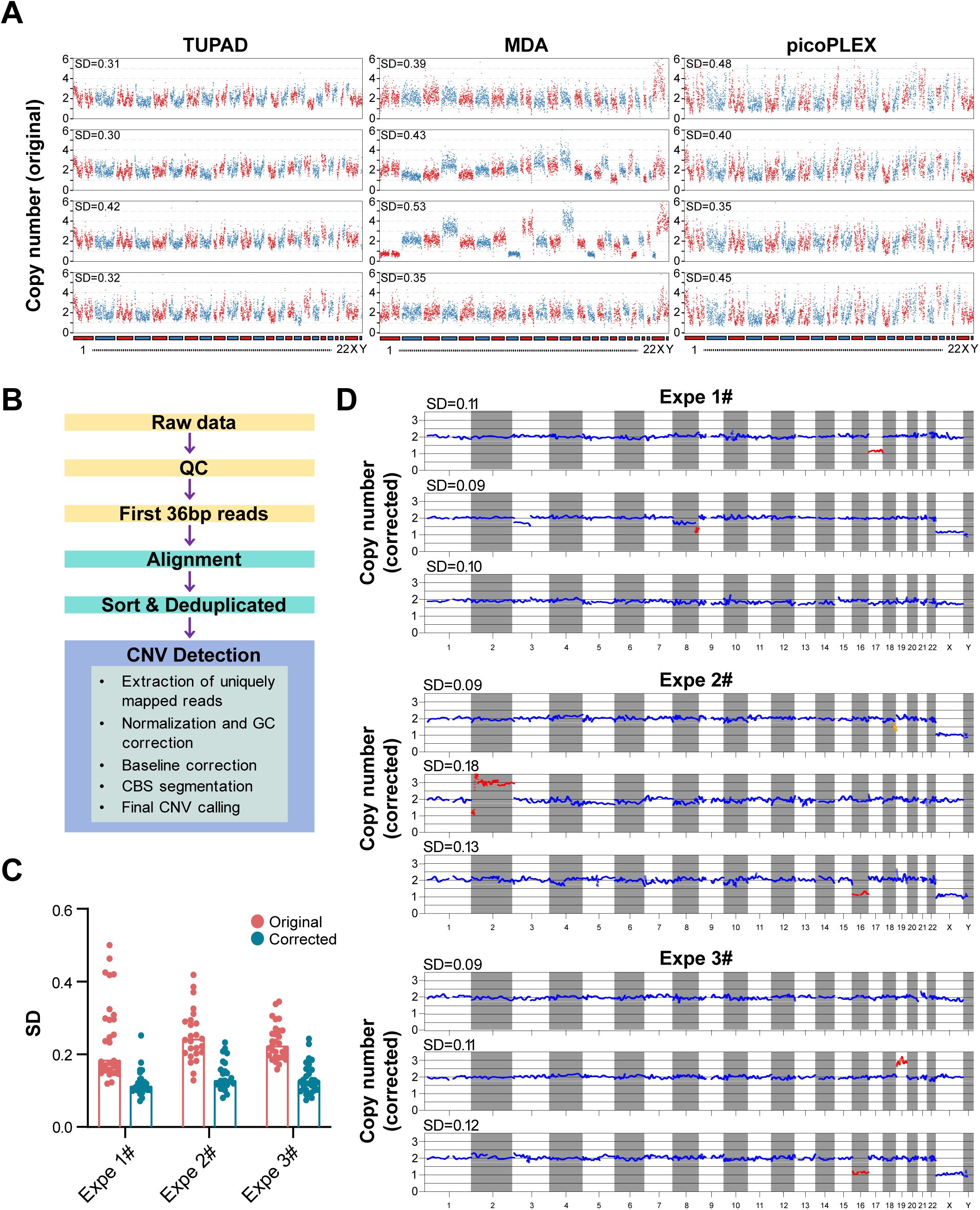
Copy-number variation (CNV) analysis of experimental embryos. (A) Original copy-number profiles of experimental embryos generated using TUPAD, MDA, and picoPLEX. The standard deviation (SD), indicated in the upper-left corner of each panel, was calculated from raw read counts within 600-kb genomic windows. (B) Schematic overview of the baseline CNV algorithm workflow. Detailed procedures are described in the Methods section. (C) Summary of SD statistics for copy-number analysis of experimental embryos generated using TUPAD after baseline algorithm correction. Expe 1# (n = 30), Expe 2# (n = 25), and Expe 3# (n = 30) correspond to three independent experimental batches. (D) Copy-number profiles of experimental embryos generated using the baseline CNV algorithm shown in panel (B). Expe 1# (n = 30), Expe 2# (n = 25), and Expe 3# (n = 30) represent three independent experimental batches as in panel (C), with three representative embryos selected from each batch for visualization.

TUPAD achieved a 100% success rate in the experimentation and sequencing of three sets of clinical embryo samples, encompassing 85 specimens total. Following algorithmic correction, uniformity was improved across all samples (Fig. 5C), enabling the detection of genuine CNV variants in the clinical embryos (Fig. 5D). Notably, TUPAD retains high coverage and uniformity in practical samples to deliver more accurate results, while offering enhanced experimental efficiency: library preparation is completed in 1.5 hours, reducing processing time by 50% compared to conventional WGA workflows.

## Discussion

In this study, we developed TUPAD, a pre-amplification-free, ultra-low input library preparation strategy that facilitates highly uniform single-cell genomic sequencing and supports concurrent analysis of multiple preimplantation genetic testing (PGT) modalities in a single assay. By circumventing pre-amplification, TUPAD directly addresses long-standing challenges in low-input sequencing, including genomic coverage limitations, amplification bias, and allelic dropout.

A key innovation driving this work was the identification of adapter dimers as the primary impurity in transposase-based low-input library construction. Similar to conventional workflows, excessive adapter dimers occupy sequencing flow cell space, invalidate library quality, and severely compromise construction success rates. To address this bottleneck, we established an adapter-dimer-free technology platform: first, we used T7 endonuclease I (T7 endo I, which specifically recognizes DNA base mismatches) combined with an optimized N001 buffer, eliminating over 90% of T4 DNA ligase-derived dimers; second, we modified primer and amplifier designs to prevent co-enrichment of transposase-intrinsic dimers during PCR, achieving complete eradication of such artifacts and drastically improving ultra-low-input library success rates.

Critical to translating this adapter-dimer-free system to single-cell applications was the deliberate selection of Tn5 transposase over T4 ligase-dependent strategies, motivated by fundamental limitations of T4 ligase in direct post-lysis library construction. T4 ligase exhibits three core constraints: (1) it requires purified, high-integrity genomic DNA templates; (2) it is highly sensitive to the complex matrix of cell lysis buffers (residual proteases, chaotropic agents, cellular metabolites), which inhibit catalytic activity and reduce ligation efficiency; (3) its incompatibility with unpurified lysates necessitates laborious DNA purification and quality control steps—impractical for single-cell samples due to inevitable nucleic acid loss and potential bias introduction. In stark contrast, Tn5 transposase functions robustly in mild lysis buffers and executes genomic fragmentation and adapter ligation in a single one-pot reaction, obviating DNA purification entirely. This intrinsic lysis buffer tolerance and one-step architecture make Tn5 uniquely suited for direct single-cell library preparation. Leveraging this advantage, we integrated a single-cell lysis module with our Tn5-based adapter-dimer-free technology to develop TUPAD, a single-cell library assembly platform compatible with low-input scenarios.

Benchmarking experiments using 1-6 cell PBMC micro-samples demonstrated that TUPAD produces highly uniform genome coverage, with coverage breadth comparable to MDA and coverage uniformity exceeding that of PicoPLEX. This uniformity enabled accurate detection of copy number alterations in PGT-A reference samples. For single-nucleotide and small indel variants, TUPAD showed improved sensitivity and fidelity relative to WGA-based methods, including reduced allelic dropout rates and allele frequency distributions that more closely matched bulk DNA references, particularly at clinically relevant loci such as BRCA. In embryo biopsy samples, TUPAD consistently maintained superior coverage uniformity for aneuploidy detection. In parallel, we established a dedicated bioinformatics pipeline and baseline calibration framework to standardize variant calling and facilitate clinical translation. The principal distinction between TUPAD and existing WGA-based approaches lies in its elimination of pre-amplification as a prerequisite for genomic enrichment. WGA methods are inherently susceptible to stochastic amplification effects, leading to uneven representation of low-copy regions, poor coverage of GC-rich or structurally complex loci, and the formation of chimeric reads. While individual strategies such as MDA, MALBAC, PicoPLEX, PTA, and LIANTI attempt to mitigate specific aspects of these limitations, each remains constrained by trade-offs between coverage breadth, uniformity, reproducibility, and workflow complexity. By directly incorporating native genomic templates into sequencing libraries, TUPAD avoids amplification-induced distortions altogether, resulting in improved inter-cell reproducibility and more reliable detection of CNVs, SNVs, and structural rearrangements within a single assay.

Beyond PGT, TUPAD is broadly applicable to other areas of single-cell and low-input genomics. Its high-fidelity and uniform coverage are well suited for cancer genomics, where accurate detection of rare somatic variants and structural alterations is essential. The streamlined workflow and compatibility with standard reaction volumes also support scalability for population-level studies and integration into automated clinical pipelines. Future efforts will focus on further increasing throughput, expanding compatibility with additional sample types, and exploring integration with other single-cell omics modalities. Potential limitations, including performance on highly degraded DNA and ultra-high throughput implementations, warrant continued optimization.

Finally, although TUPAD eliminates amplification-induced bias at the genomic enrichment stage, downstream library amplification remains necessary to generate sufficient material for sequencing. Because this amplification occurs after uniform library construction, it introduces substantially less distortion than WGA.

In summary, TUPAD establishes a robust framework for pre-amplification-free single-cell library preparation. By combining Tn5-mediated direct library construction with comprehensive suppression of adapter-dimer artifacts, it overcomes key biochemical and technical barriers that have constrained low-input sequencing. These features position TUPAD as a practical and reliable solution for integrated PGT analysis and broader single-cell genomic applications.

## Author contributions

W,Y.;G,J;C,L;S,T;Q,Z;Z,L;conceived the project and designed the experiments.W.k.;G,H.W,K,N,R; performed .bioinformatics data analysis;W,M;Z,H;W,H;L,Y; A,Y; O,Y;C,R;implementation of wet experiments;T,Q;W,Y;G,J;W,K;Z,X; wrote the manuscript with input from all other authors;X,Y;prepared the primate samples; All authors edited and proofread the manuscript.

## Materials and Methods

### Standard Reference gDNA

Standard reference gDNA used in low input DNA Library experiment was purchased from GeneWell Biotechnology Co., Ltd. (Guangdong, China). Including BRCA gDNA reference standard (#CA3387), pathogenic copy number variation gDNA reference standard and autosomal trisomy gDNA reference standard. Genomic DNA from HG001 (NA12878) was obtained from the Coriell Institute.

### Human Peripheral Blood Mononuclear Cell (PBMC)

EDTA-anticoagulated peripheral whole blood samples were obtained from normal adult human donor. incubated immediately after collection at room temperature (20-25℃), with gentle end-to-end shaking for 2 to 48h. The PBMC fractions were isolated usingthe Ficoll. Whole blood samples were pre-diluted 1:1 with phosphate-buffered saline (PBS) without Ca2+and Mg2+(Gibco, Cat# J67802-AP) and layered carefully on an equal volume of Ficoll-Paque PREMIUM (Cytiva, Cat# GE17-5442-03). The PBMC fraction was isolated following centrifugation at 400×g for 30 min at room temperature with the brake off. Single PBMC cell was picked with capillary pipette, PBMC was diluted with PBS, single picking was micromanipulation under microscope. PBMC genomic DNA isolation using the FastPure Blood/Cell/Tissue/Bacteria DNA Isolation Mini Kit (Vazyme, Cat# DC112).

### Researh embryo experiment

Researh embryo sample experiment was performed at the first affiliated hospital of Sun Yat-sen university under the IEC for clinial research and animal trial approval (Ethical Approval Number: [2021]561).

### Library construction

Genomic DNA (gDNA) was quantified using the Equalbit 1 × dsDNA HS Assay Kit (Vazyme, Cat#EQ121). DNA library was constructed with fragmentation library preparation kit (Vazyme, Cat#UND637) or transposase library preparation kit (Vazyme, Cat#TD503, Cat#UTD523). The entire library preparation procedure was performed following the manufacturer’s instructions. Purification of samples was performed using an identical volume of VAHTS DNA Clean Beads (Vazyme, Cat#N411), according to the manufacturer’s instructions. Quality control was performed using an Agilent 2100 Bioanalyzer and DNA 1000 kit.

### Adapter-free Library construction

In conventional enzymatic fragmentation and ligation-based library preparations, adapter dimers were actively removed during the bead purification step. Specifically, a dimer-wash buffer (Vazyme, Cat#N001) was applied during magnetic bead purification to efficiently remove short, self-ligated adapter dimer. Additionally, T7 endonuclease I (Vazyme, Cat#EN303) was utilized to cleave the self-ligated adapter dimer at T:T mismatch.

For transposase-based library preparation (tagmentation), a unique approach was employed to actively block the amplification of adapter dimers generated during the reaction. This enables robust, adapter dimer-free library construction even with low starting material input, this strategy has been applied in the reagent UTD523.

### Single cell whole genome amplification

For conventional single cell library construction, single cell or low-input cell first was performed whole genome amplification (WGA) by MDA (Qiagen, 150345), MALBAC (Yikon, Cat#KT110700110), picoPLEX (Takara, Cat# R300718). The library was prepared by transposase-based library preparation kit (Vazyme, UTD523)

### TUPAD

Single-cell Transposase-based Adapter-Free Library assembly Enabling Clinical-application TUPAD methods has been developed standard solution kit (Vazyme, Cat#SC701). For single-cell and low-input cells (1 to 10) were first lysed to release genomic DNA. Subsequently, the released gDNA was subjected to tagmentation using the Tn5 transposase, which simultaneously fragments the DNA and integrates specific sequences (tags) onto the ends of the fragments. The final sequencing library was then generated via Adapter-free amplification step, utilizing primers complementary to the Tn5-integrated tags.

### Preparation of Artificial Adapter-Dimer substrates

The Artificial Illumina Truseq Y-shape Adapter-Dimer substrates were obtained by using fragmentation library preparation kit (Vazyme, Cat#UND637). Under no input gDNA, only Truseq Y-shape Adapter were self-ligated adapter-dimer, then adapter-dimer were purified with VAHTS DNA Clean Beads (Vazyme, Cat#N411). According to the experimental requirements, the different rate of adapter-dimer was added into libraries in the following proportions: 1:5, 1:10, 1:20, 1:50 respectively.

### Eliminate the adapter dimer produced by Tn5 transposase-based library construction

The first method involves using 3’ end elongation primers to replace the original amplification primers.

The second method involves performing 1 to 3 thioester modifications at the 3’ ends of the primers that are extended at the 3’ end.

The third method involves replacing the original amplification mix in the kit with 2 × KeyPo SE Master Mix (Dye Plus) (Vazyme PK512).

### DNA sequencing

All libraries were sequenced on an Illumina HiSeq 2000 according to standard procedures.

### Sequencing Data Processing and Benchmarking

Adapter sequences and low-quality bases at read termini were removed using Cutadapt(v1.18)^29^. The trimmed reads were aligned to the hg38 human reference genome using bwa-mem2 (v2.2.1)[10.1109/IPDPS.2019.00041] with default parameters. PCR duplicates were subsequently marked and removed using Picard (v2.18.29) [https://github.com/broadinstitute/picard], thereby ensuring comparable effective sequencing depth across samples and minimizing amplification-related biases.

To enable fair benchmarking across datasets, all BAM files were downsampled to predefined read counts using Samtools (v1.5)^30^. Alignment and sequencing quality metrics, including mapping rate and insert size distribution, were collected from the final BAM files using Qualimap(2.2.2a)^31^.

Genome-wide coverage statistics were calculated using mosdepth(v0.3.3)^32^. Reads with low mapping quality were excluded prior to coverage calculation, and genome coverage was assessed at 1× depth as well as across multiple depth thresholds to evaluate coverage breadth and depth distribution. Coverage uniformity was further assessed by generating Lorenz curves using Picard CollectWgsMetrics. All downstream statistical analyses and visualizations were performed using custom R scripts.

### Uniformity characterization

Coverage uniformity was evaluated across the genome using different bin sizes generated with bedtools makewindows. To avoid biases introduced by poorly mappable regions, bins overlapping ENCODE^33^ blacklist regions and UCSC^34^ gap regions were excluded from downstream analyses. Read counts for each remaining bin were calculated using Sambamba(v0.6.6)^35^, and the resulting per-bin coverage profiles were used to compute coverage uniformity metrics. The coefficient of variation (CV) was calculated for each bin size to quantify global coverage dispersion across the genome. CV was defined as:

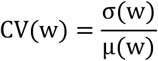

where σ(w) and μ(w) represent the standard deviation and mean coverage, respectively, of genomic bins with window size w. In addition, coverage smoothness was assessed using the median absolute pairwise difference (MAPD), which captures local coverage fluctuation between adjacent genomic bins. MAPD was calculated as:

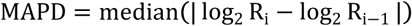

where R_i_ and R_i−1_ denote the coverage of adjacent bins along the genome. Lower CV and MAPD values indicate improved coverage uniformity and reduced local amplification bias.

### SNV and INDEL Detection and Analysis

For BRCA mutation detection, aligned reads were first processed to remove PCR duplicates using Picard MarkDuplicates. To enable fair comparison of variant detection performance across different amplification strategies, all WGA-derived datasets were downsampled to a uniform sequencing depth of 15×, together with TUPAD and WGS inputs.

Single-nucleotide variants (SNVs) and short insertions and deletions (INDELs) were called using GATK4 HaplotypeCaller in GVCF mode for each sample, followed by joint genotyping using GenotypeGVCFs to generate a unified VCF across samples.A truth set of heterozygous SNVs and INDELs was defined based on the bulk WGS replicates. Variants were considered true heterozygous sites if they satisfied a minimum total read depth threshold in all WGS replicates and showed consistent genotypes as determined by HaplotypeCaller.

For TUPAD and WGA samples, variants with a read depth greater than 10× were evaluated against the truth set. A variant was classified as a true positive (TP) if its genotype matched the truth set; otherwise, it was classified as a false positive (FP). Truth-set variants that were not detected in the corresponding single-cell or WGA samples were classified as false negatives (FN). Variant detection performance was quantified using precision, sensitivity (recall), and F1 score, defined as:

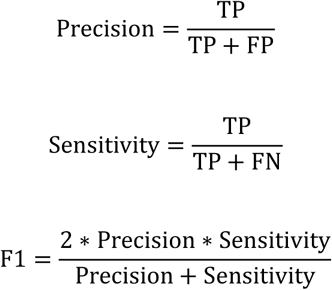

Allelic imbalance was further assessed by quantifying allele dropout (ADO) events. A non-reference allele dropout (NR-ADO) event was defined as a heterozygous variant in the WGS truth set for which no reads supporting the non-reference allele were observed in the corresponding single-cell or WGA sample. The NR-ADO rate was calculated as the proportion of NR-ADO events among all heterozygous variants in the truth set. Similarly, reference allele dropout (Ref-ADO) was defined and calculated when the reference allele was absent at heterozygous sites.

### Copy Number Variation (CNV) Analysis

CNV analysis was performed using a reference-based calibration pipeline optimized for high-sensitivity detection. Raw sequencing reads were quality-filtered and trimmed to retain the first 36 bp, then aligned to the human reference genome (hg19) using BWA (v0.7.13)^36^Alignments were sorted and PCR duplicates removed prior to downstream analysis. The genome was partitioned into non-overlapping 600-kb bins, with bin sizes selected to ensure ≥50 unique reads per bin at ∼0.3 million unique reads. Unique read counts for each bin were normalized, and outlier bins with counts outside 0.7 – 1.3 × the initial autosomal mean were excluded.

To correct for GC-content–related coverage bias, a LOESS (locally estimated scatterplot smoothing) regression was fitted between bam_cnt and the GC content of each bin. The fitted model was used to predict Z-score values for GC normalization. GC-normalized bin counts (gc_cnt) were further corrected using a baseline reference file, which contains the median and standard deviation (SD) of gc_cnt values derived from reference samples for each bin. The corrected gc_cnt values were computed using a predefined normalization formula based on baseline statistics.To prevent inflation of normalized values caused by extremely low read counts, any gc_cnt value below 0.1 was set to 0.1 and excluded from the normalization calculation. For each genomic bin i of the testing sample, if 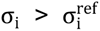, the raw read count C_i_ was normalized according to:

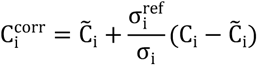

where C̃_i_ and σ_i_ denote the median and standard deviation of GC-content–normalized read counts across all baseline samples for bin i, respectively, and 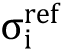 represents the median reference standard deviation estimated from baseline samples. 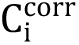 is the final GC-corrected read count of the testing sample for bin i.

Final bin weights were assigned based on variability observed in the baseline file. Specifically, if a bin exhibited a baseline SD > 1.0, its weight was set to the inverse of its variance; otherwise, the bin weight was set to 1. The normalized log2 ratio values were segmented using the Circular Binary Segmentation (CBS) algorithm^37^. Segment-level log2 ratios were transformed to linear copy number ratios using:

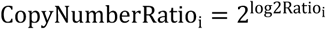

Mosaic CNVs were determined by recalculating segment means after excluding outlier bins. Final CNV calls were reported for segments with estimated mosaic ratios between 30% and 70%.

---

**Figure S1.**
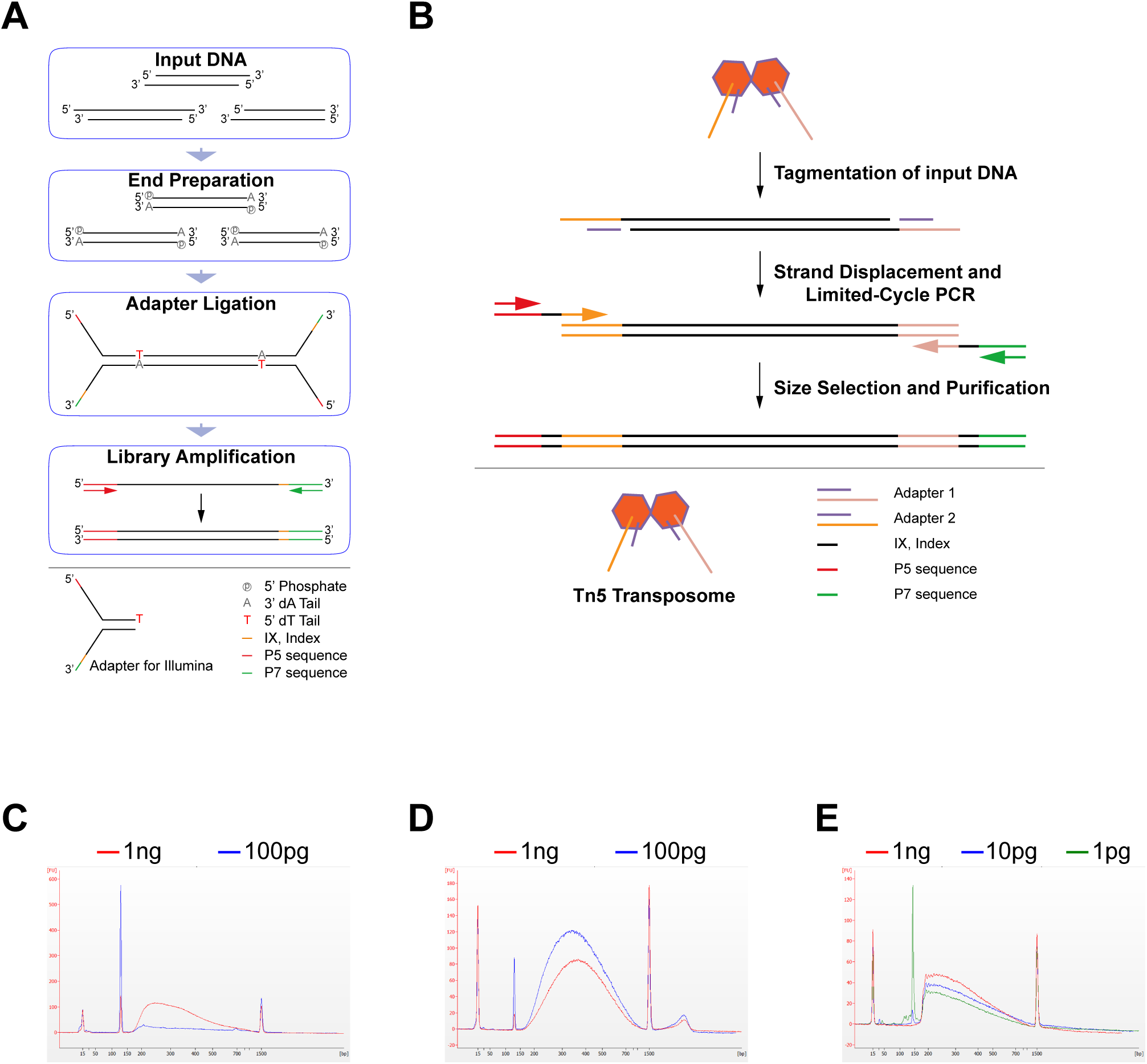
DNA library preparation workflows and input-dependent library size distributions. (A) Schematic overview of the Illumina ligation-based library preparation workflow, in which genomic DNA is fragmented by ultrasonication or enzymatic digestion, followed by end repair and adapter ligation to generate sequencing libraries. (B) Schematic overview of the transposase-based library preparation workflow, where the Tn5 transposome complex simultaneously fragments DNA and inserts sequencing adapters, with the final library structure completed through PCR amplification. (C) Bioanalyzer electropherograms of DNA libraries generated by ultrasonication-based fragmentation using different input amounts of Staphylococcus aureus DNA. The red trace represents 1 ng, and the blue trace represents 100 pg input. (D) Bioanalyzer electropherograms of DNA libraries generated by the enzymatic fragmentation method using different input amounts of Staphylococcus aureus DNA. The red trace represents 1 ng, and the blue trace represents 100 pg input. (E) Bioanalyzer electropherograms of DNA libraries generated by the transposase-based library preparation method using different input amounts of HEK293T cell DNA. The red trace represents 1 ng, the blue trace represents 10 pg, and the green trace represents 1 pg input.

**Figure S2.**
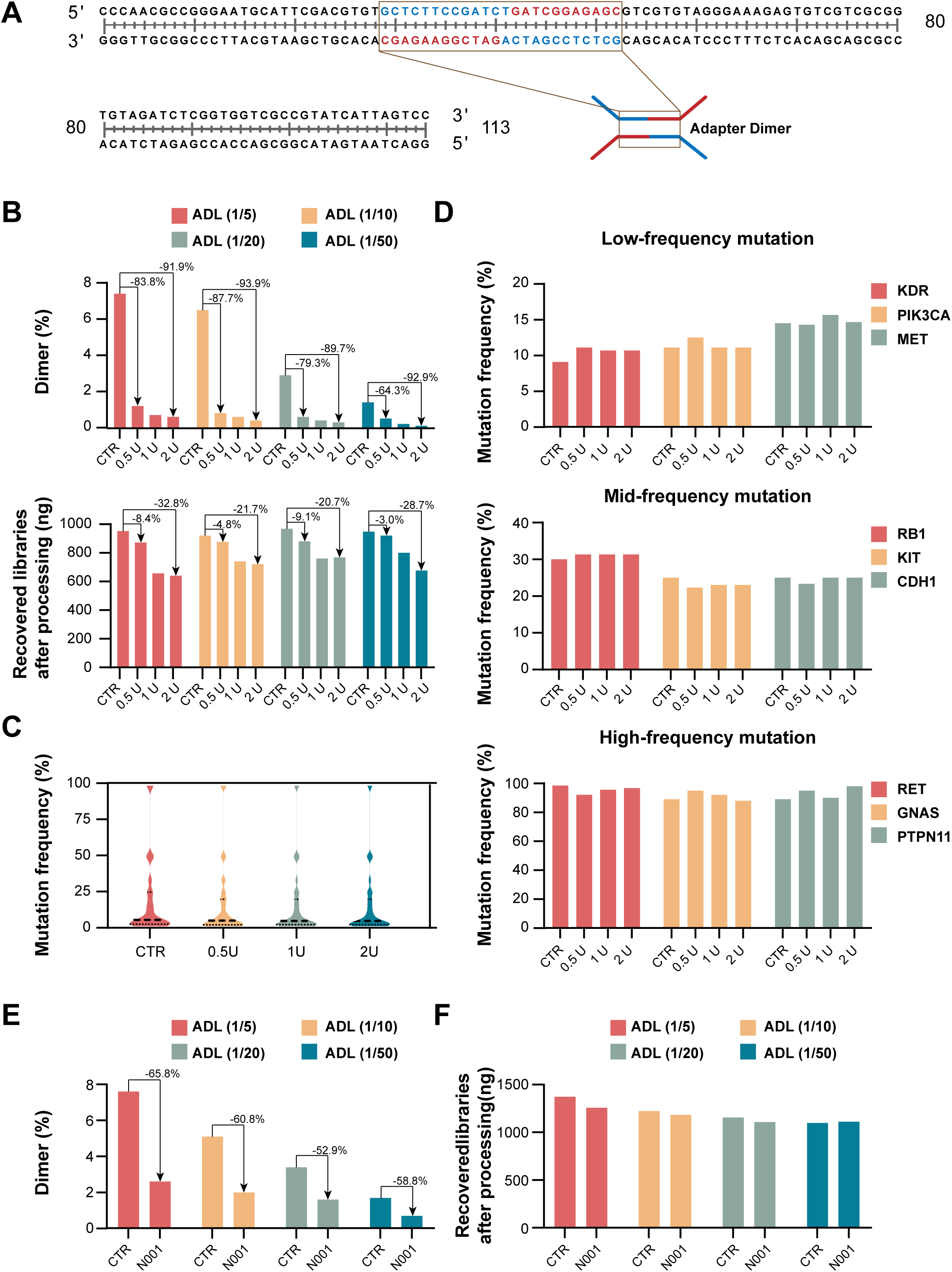
Elimination of adapter dimer in ligation-based low-input library preparation. (A) Sanger sequencing validation of Illumina Y-adapter–derived dimer sequences. The brown boxes highlight the complementary regions responsible for adapter-dimer formation. (B) Adapter-dimer library (ADL) model and T7 Endonuclease I treatment. The ADL model was constructed by mixing adapter dimers with a normal library at different mass ratios (adapter-dimer to normal library ratios of 1:5, 1:10, 1:20, and 1:50; color-coded as red, yellow, green, and blue, respectively). The upper panel shows the remaining proportion of adapter dimers after treatment with increasing amounts of T7 Endonuclease I, and the lower panel shows the corresponding library yield. CTR represents the control group, while 0.5U, 1U, and 2U represent the corresponding T7 Endonuclease I treatment groups. The arrows indicate the percentage reduction compared to the control after the treatment. (C) Statistical analysis of the detected mutation frequency following treatment with varying amounts of T7 Endonuclease I. CTR represents the control group, while 0.5U, 1U, and 2U represent the corresponding T7 Endonuclease I treatment groups. (D) Mutation frequency distributions stratified by low-, medium-, and high-frequency variants. The top, middle, and bottom panels show mutation frequencies detected in genes carrying low-, medium-, and high-frequency mutations, respectively, after treatment with varying amounts of T7 Endonuclease I. (E) Adapter-dimer rate of the ADL model after treatment with a magnetic bead elution buffer (N001). CTR represents the control group, and the arrows indicate the percentage reduction compared to the control after the N001 buffer treatment. (F) Library yield of the ADL model after treatment with a magnetic bead elution buffer (N001).

**Figure S3.**
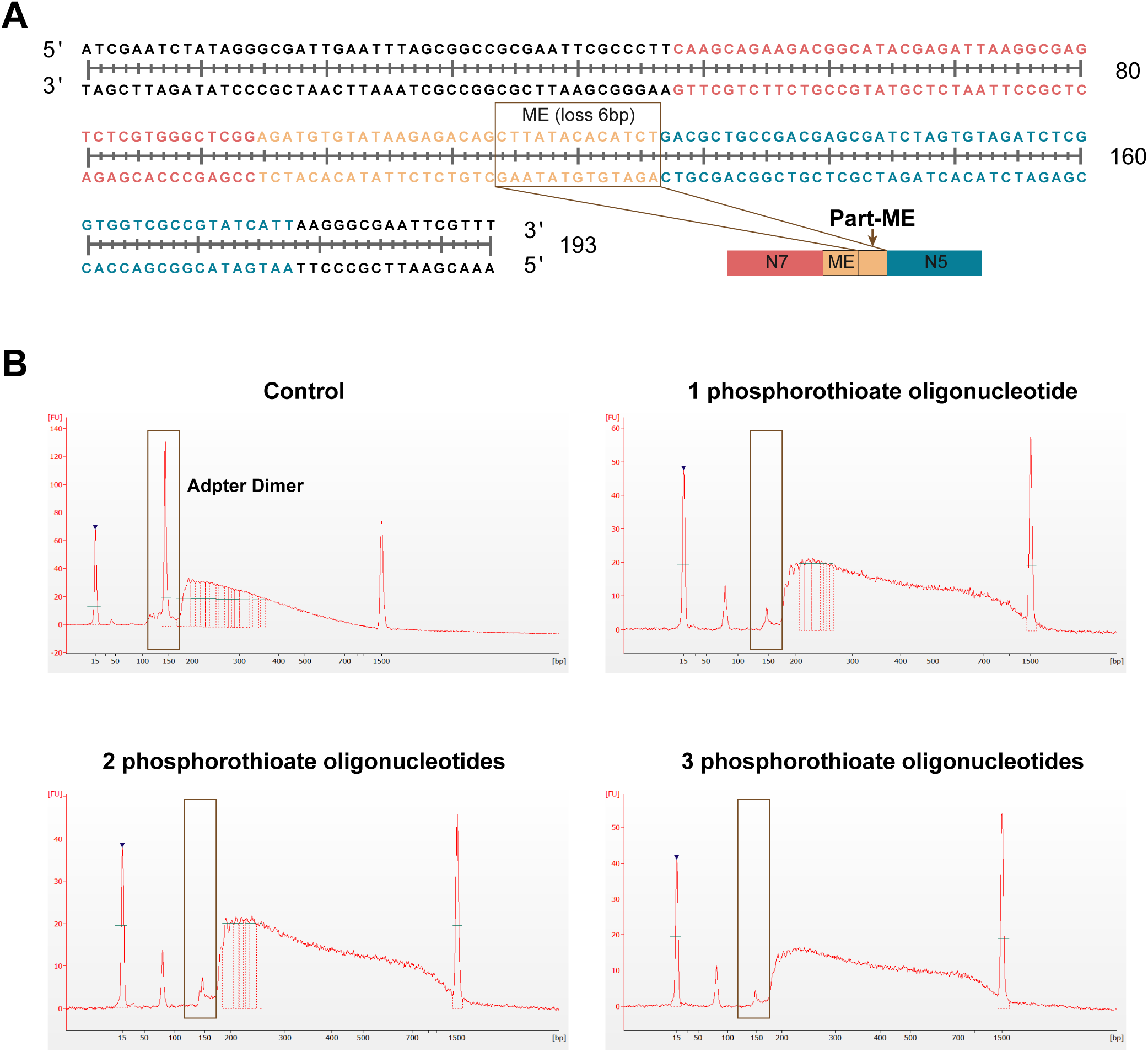
Elimination of adapter dimers in Tn5-mediated low-input library preparation. (A)Sequencing validation of library structures containing part-ME sequences. The brown box highlights part-ME sequences with a 6-bp 5’ truncation, while the overall library structure retains a normal N5 index, complete ME sequence, truncated ME sequence, and N7 index in order. (B) Size distribution of libraries analyzed by Agilent 2100 Bioanalyzer. Control represent library amplified with conventional index primers, 1-3 phosphorothioate oligonucleotide represent libraries amplified with different numbers of phosphorothioate modifications in the end of extended index primers. The black boxes highlight the regions responsible for adapter-dimer formation.

**Fig S4.**
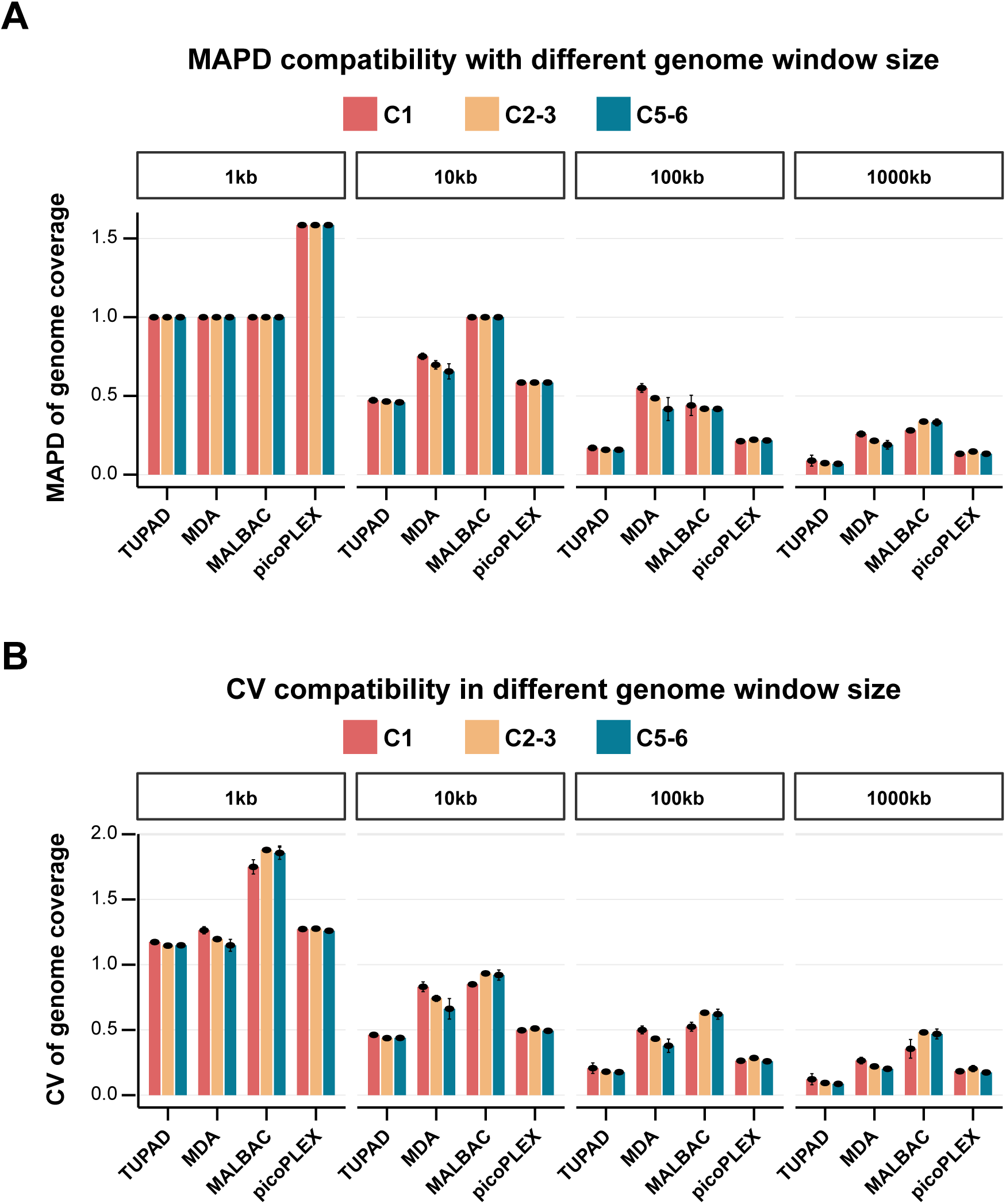
Genome coverage uniformity across different input cell numbers. (A) Genome coverage uniformity measured by the MAPD across varying genomic window sizes and increasing input cell numbers (1 cell, 2-3 cells, and 5-6 cells), with two technical replicates per method. TUPAD consistently exhibited lower MAPD than the other three WGA methods for window sizes greater than 10 kb. (B) Genome coverage uniformity measured by the CV across different genomic window sizes and increasing input cell numbers (1 cell, 2-3 cells, and 5-6 cells), with two technical replicates per method. TUPAD showed lower CV values than the other three WGA methods across all window sizes.

**Figure S5.**
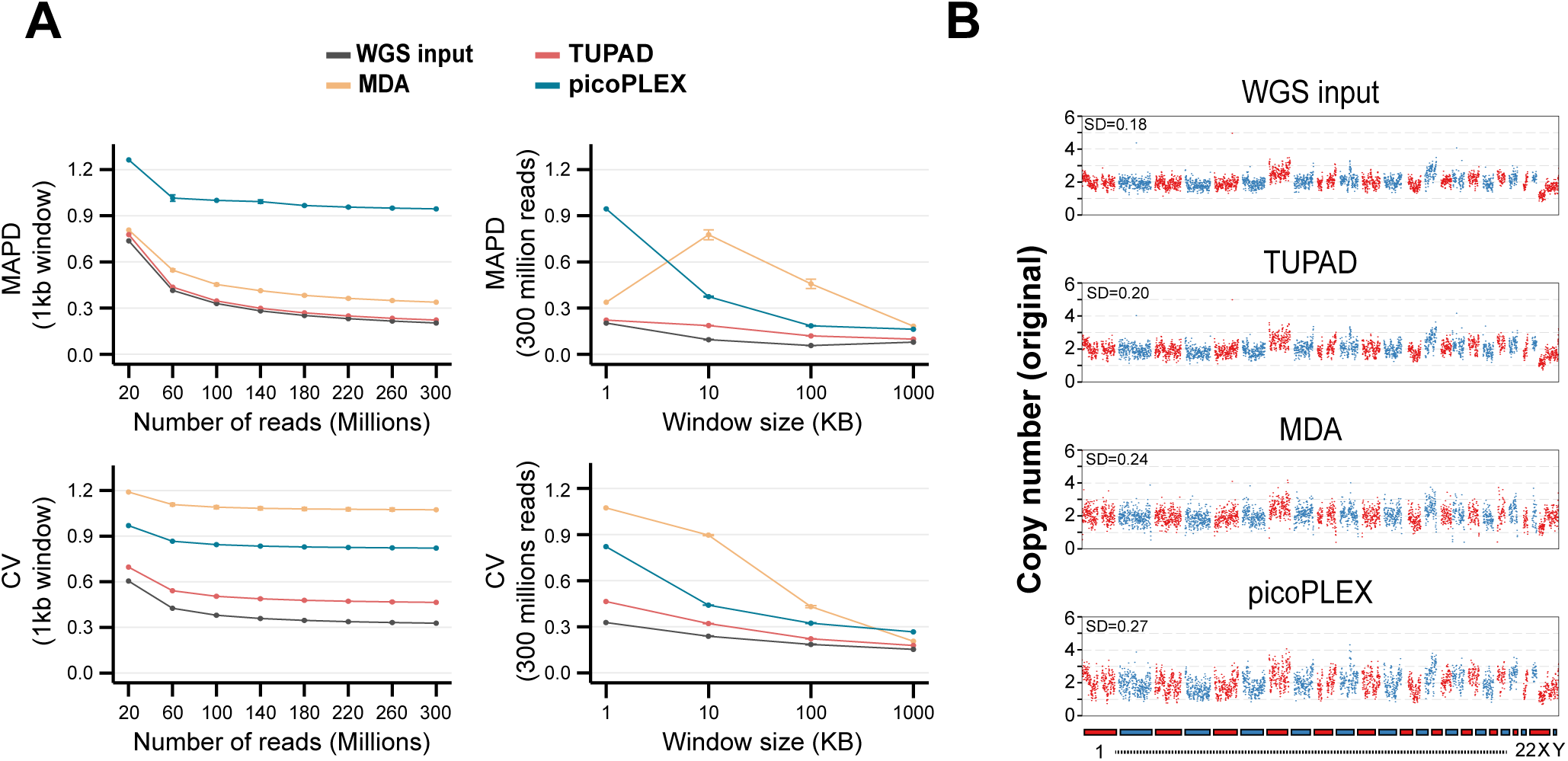
Genome coverage uniformity and copy-number analysis of BRCA reference standards. (A) Genome coverage uniformity measured by the median absolute pairwise difference (MAPD) with increasing numbers of single-end sequencing reads using 1 kb genomic windows. (B) Genome coverage uniformity measured by MAPD across window sizes ranging from 1 kb to 1000 kb using 300 million single-end reads (approximately 15× whole-genome coverage). (C) Genome coverage uniformity measured by the coefficient of variation (CV) with increasing numbers of single-end sequencing reads using 1 kb genomic windows. (D) Genome coverage uniformity measured by CV across window sizes ranging from 1 kb to 1000 kb using 300 million single-end reads. (E) Original copy-number profiles across all 23 chromosomes of BRCA reference standards generated using WGS input, TUPAD, MDA, and picoPLEX.

**Figure S6.**
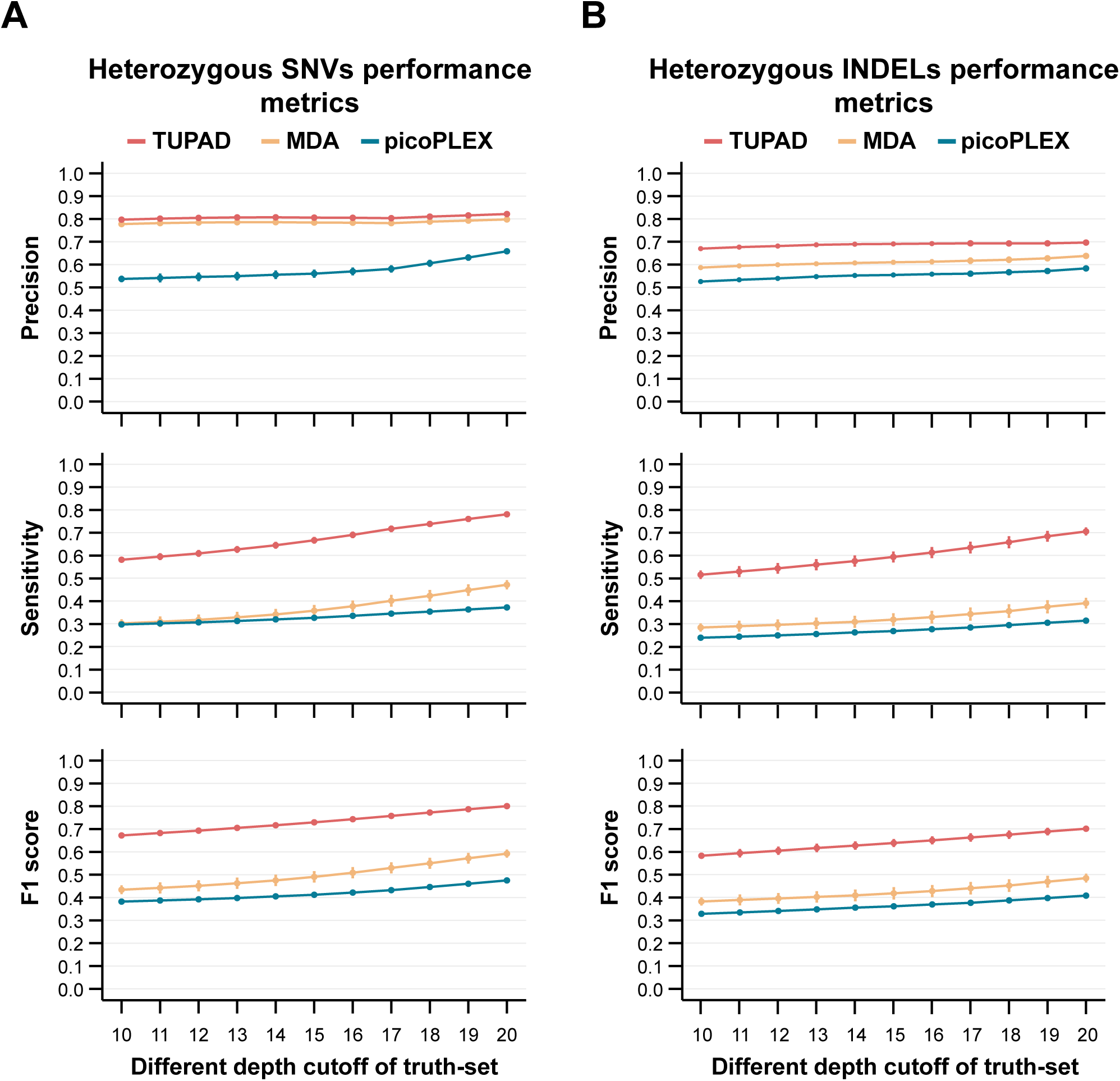
Variant detection performance of standard BRCA gDNA reference. (A) Comparison of heterozygous SNV detection performance across variable truth sets defined by different read-depth cutoffs. True heterozygous SNVs were defined as sites supported by at least the cutoff depth in both WGS input replicates, generating different truth sets at each cutoff. Detection performance of each method was evaluated using precision, sensitivity, and F1 score. TUPAD maintained consistently high precision across all depth thresholds. (B) Comparison of heterozygous INDEL detection performance using the same depth-defined truth sets. Performance was evaluated with precision, sensitivity, and F1 score, with TUPAD consistently achieving the highest precision.

**Figure S7.**
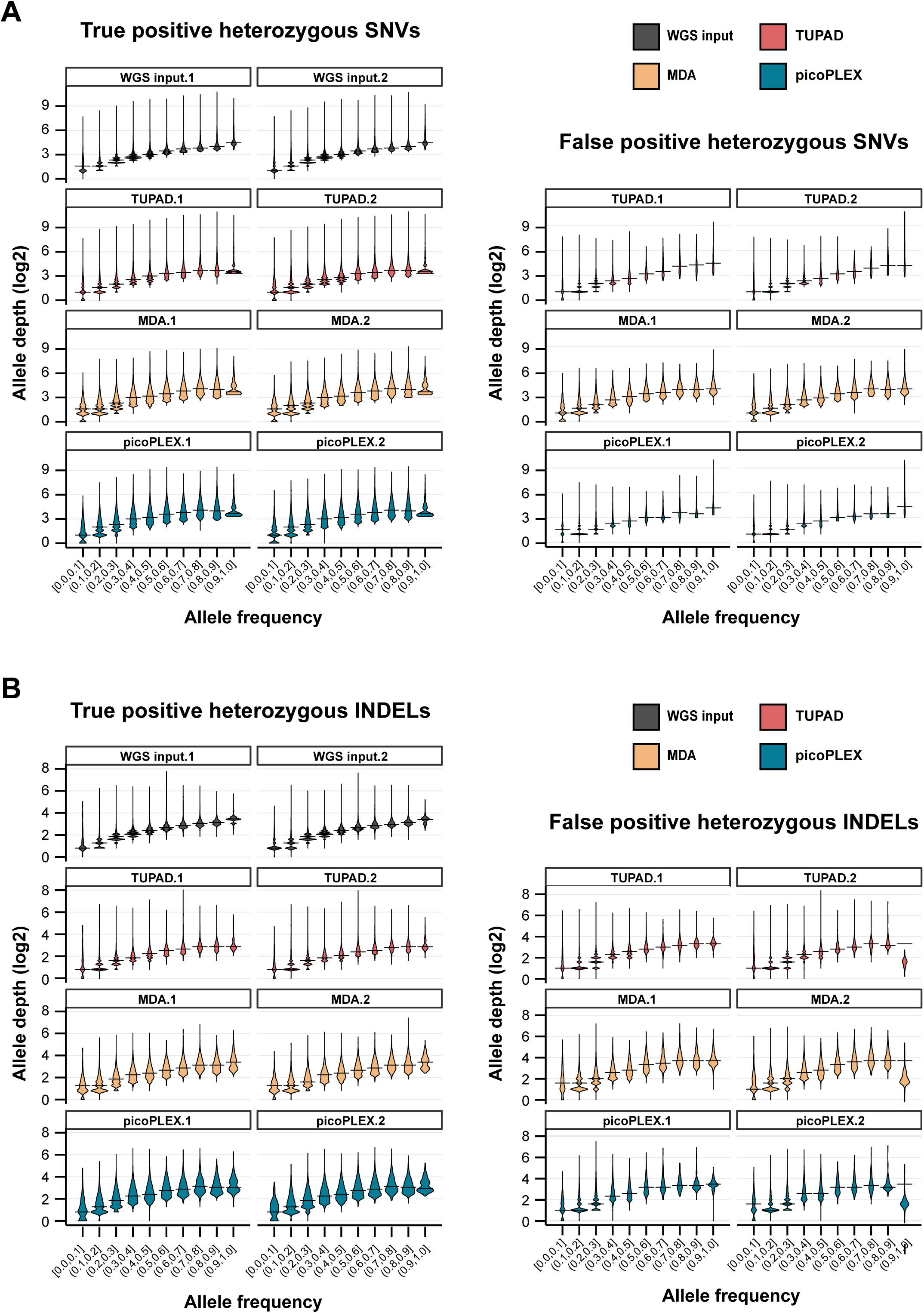
Distribution of SNVs and INDELs across allele-frequency groups. (A) Violin plots showing read-depth distributions of heterozygous SNVs across allele-frequency groups for each method. Left and right panels show false positive (FP) and true positive (TP) SNVs, respectively. The truth set was defined using a read-depth threshold of >= 15 reads in both WGS input samples. TP and FP definitions are detailed in the Methods. Overall, TUPAD’s SNV distribution closely resembles that of the bulk WGS input. (B) Violin plots showing read-depth distributions of heterozygous INDELs across allele-frequency groups for each method, analyzed using the same pipeline as SNVs (panel A).

**Figure S8.**
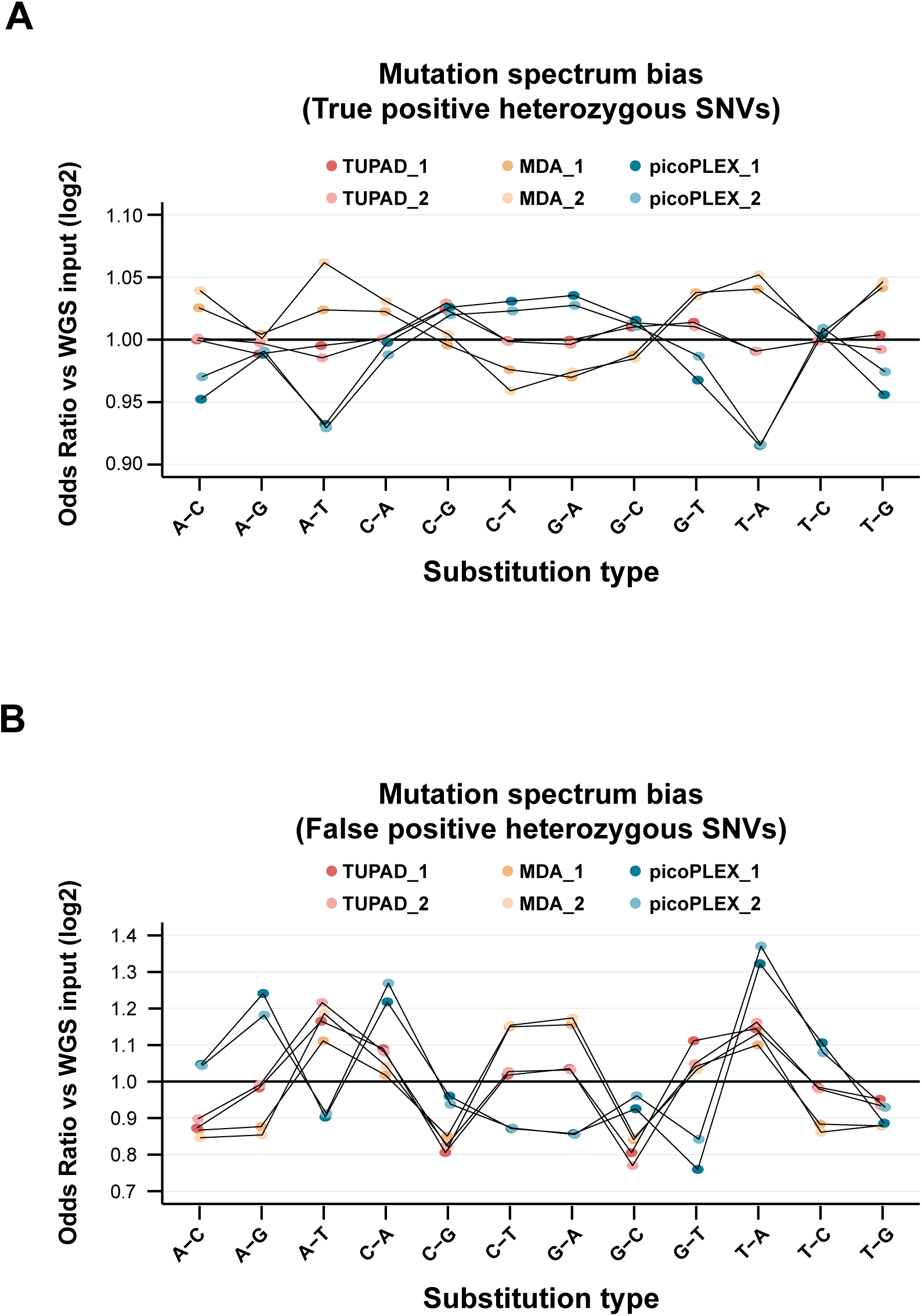
Mutation spectrum bias of SNVs. (A) Line plots showing mutation spectrum bias of true positive (TP) heterozygous SNVs across 12 base-substitution types for each method. For each method, the odds ratio was calculated by comparing the proportion of each substitution type to that in the WGS input, followed by log_2_ transformation. (B) Line plots showing mutation spectrum bias of false positive (FP) heterozygous SNVs across the same 12 substitution types. Odds ratios were calculated using the same approach as in panel (A).

